# Competition within species determines the value of a mutualism between species

**DOI:** 10.1101/402271

**Authors:** Syuan-Jyun Sun, Nicholas P.C. Horrocks, Rebecca M. Kilner

## Abstract

Social interactions within species, and mutualisms between species are both well characterised, but their influence on each other is poorly understood. We determined how interactions among burying beetles *Nicrophorus vespilloides* influence the value of their interactions with the mite *Poecilochirus carabi.* Beetles transport these mites to carrion, upon which both species breed. We show that mites help beetles win intraspecific contests for this scarce resource: mites raise beetle body temperature, which enhances beetle competitive prowess. However, mites confer this benefit only upon smaller beetles, which are otherwise doomed by their size to lose contests for carrion. Larger beetles need no assistance to win a carcass and lose reproductive success when breeding alongside mites. We conclude that social interactions within species explain whether interactions with another species are mutualistic or parasitic.

**One Sentence Summary:** Social interactions within species can explain whether interactions with a second species are mutualistic or parasitic.

## Main Text

Mutualisms arise when two species interact cooperatively (*1*). They play a key role in contributing to biodiversity and ecosystem function (*2, 3*), and are an important source of evolutionary innovation (*1*). Nevertheless, mutualisms often degrade, both on an evolutionary timescale and within populations (*1*–*4*), yielding a spectrum of possible outcomes from mutualism to parasitism. Yet the underlying causes of this variation are poorly understood (*5*). Determining why the extent of mutualism varies among individuals is an important step in predicting the evolutionary stability of mutualisms in the longer term (*6*).

Here we propose and test the suggestion that social interactions within species contribute to variation in the outcome of a mutualism between species. Social interactions within species are important in this regard because they establish variation in the fitness benefits that might be gained from interacting with a second species. Conflict with rivals (*7*), between the sexes (*8*), or between parents and offspring (*9*), means that some individuals consistently lose fitness after a social interaction. Even when individuals cooperate, the fitness benefits are seldom distributed equally between the social partners (*10*). Social interactions with conspecifics thus place some individuals at a sustained fitness disadvantage, which potentially can be compensated through a mutualistic interaction with a second species. Mutualisms could arise if a second species directly induces a more favourable outcome from interactions with conspecifics (*11*); or if it provides a service that compensates any fitness lost to conspecifics (*12*). Either way, the outcome of social interactions with conspecifics potentially explains variation in the extent of mutualism between species: those systematically disadvantaged by their conspecifics can potentially gain more from interacting with a second species.

We tested this idea by studying the burying beetle *Nicrophorus vespilloides* and its phoretic mite *Poecilochirus carabi.* Burying beetles (*Nicrophorus* spp.) use small vertebrate carrion as a resource for breeding, and there is competition for carcasses both within and among species. The ability to secure a carcass resource strongly determines a burying beetle’s reproductive success (*13*). This is particularly the case for females who, unlike males, are likely only able to breed once or twice at most, and cannot breed successfully without a carcass. Competition among *Nicrophorus* spp. is sufficiently intense that it has likely caused character displacement, resulting in partitioning of the carrion niche among *Nicrophorus* spp (*14*). Nevertheless, individuals still face density-dependent competition for a carcass from rival conspecifics (*13*). Beetles gain ownership of a carcass by wrestling, biting and chasing competitors away (movie S1; see also (*15*)). Contests take place within each sex, and they are most likely to be won by larger beetles (*16*). Contests within *N. vespilloides* thus magnify individual size-related variation in reproductive success (*17, 18*), creating ‘winners’ who are assured a high level of reproductive success after securing a carrion breeding resource, and ‘losers’ who are much less likely to gain any reproductive success at all (*19*).

We determined whether this socially induced variation in prospective fitness influenced the extent of mutualism between *N. vespilloides* and the mite *Poecilochirus carabi* (see Supplementary Materials). These mites feed and reproduce on carrion, just like burying beetles. However, whereas burying beetles can fly and search for small dead vertebrates, mites rely entirely on their beetle hosts to transport them to a carcass (Fig. S1). There they breed alongside the beetle, using the same carrion resources for reproduction. Mites seemingly inflict no harm on their hosts while they are passengers on the beetle, which is why they are described as ‘phoretic’. During reproduction on the carrion, however, beetle-mite interactions vary from mutualism to parasitism, depending on which family member’s fitness is analysed (*11*) and on ecological factors such as the size of the carcass (*11*) and the presence of rival blowflies (*20*). Previous studies have focused entirely on interactions between mites and beetles after a carcass has been secured. We investigated whether mites could assist burying beetles in obtaining a carcass for both species to breed upon, by enhancing the beetle’s competitive prowess.

We began by staging contests between rival female burying beetles for a carcass, loading one female with 30 mites while leaving her rival mite-free (see Supplementary Materials). Female beetles were closely matched in body size so that we could attribute any variation in the outcome of a contest to the mites alone. At the start of each trial, females were placed simultaneously on a small mouse carcass and their interactions were filmed (movie S1). We found that females bearing mites were three times more likely to exhibit acts of aggression than beetles without mites (62 out of 80 aggressive behaviours recorded across all contests were initiated by beetles with mites; GLMM, χ^2^ = 21.10, d.f. = 1, *P* < 0.001). Furthermore, beetles with mites were also more likely to win contests over breeding carcasses than by chance (prop.test, *P =* 0.037; Fig. 1).

**Fig. 1.**
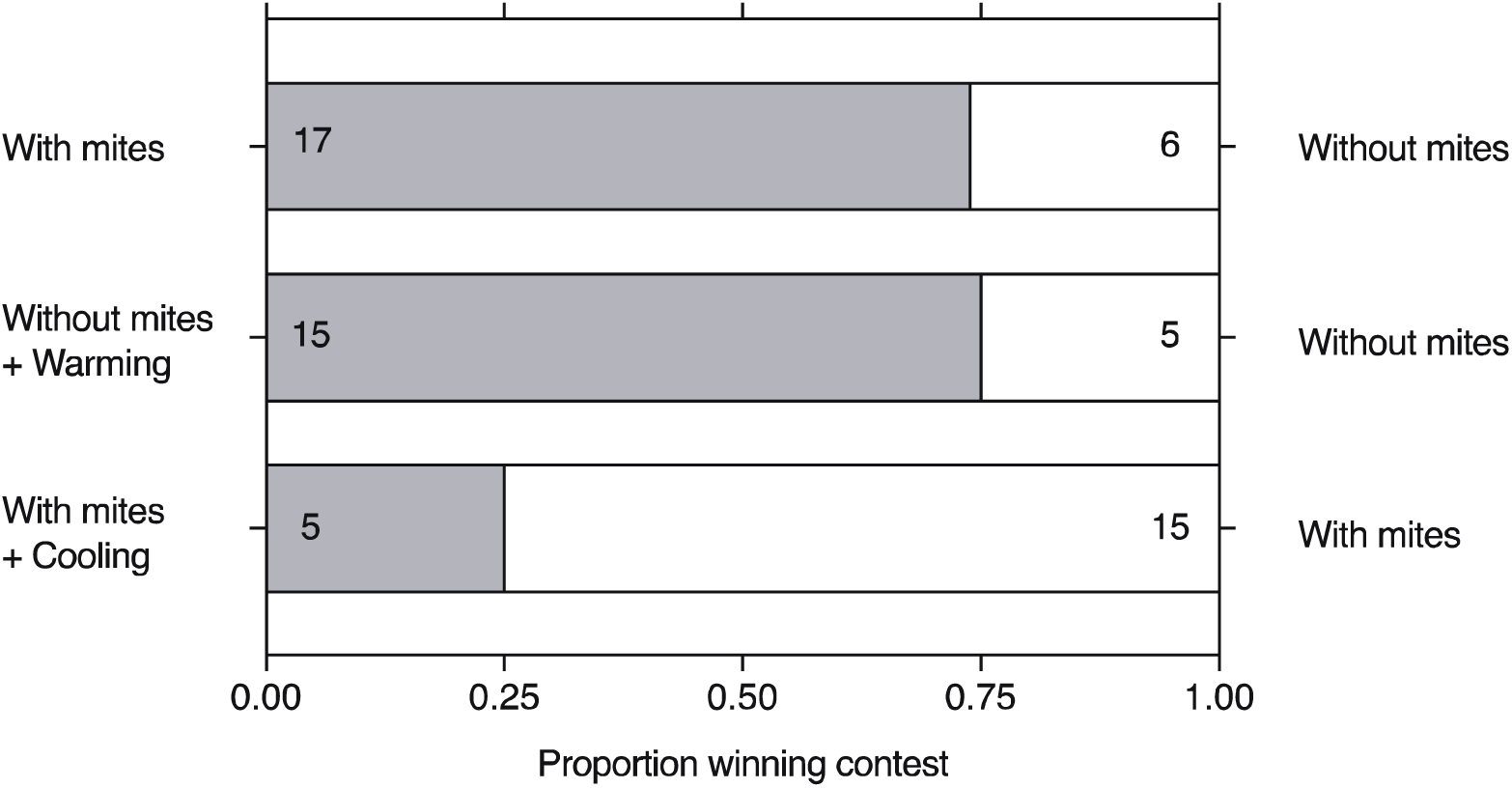
The independent effects of mites and temperature on contest outcome. Beetles were evenly matched for body size in all contests. Numbers indicate trials won by beetles.

We investigated whether the competitive superiority conferred by mites was associated with an elevated body temperature in the beetles, since this has been shown to influence competitive ability in other insects (e.g. *21*). Infrared thermography revealed that beetles bearing mites had a higher body temperature during acts of aggression than mite-free beetles (GLMM, χ^2^ = 9.54, d.f. = 1, *P =* 0.002; Fig. 2A and movie S1). Furthermore, beetles with mites had a significantly raised body temperature after fighting (GLMM, χ^2^ = 8.55, d.f. = 1, *P =* 0.003; Fig. 2B) whereas beetles without mites did not (GLMM, χ^2^ = 1.71, d.f. = 1, *P =* 0.191; Fig. 2B).

**Fig. 2.**
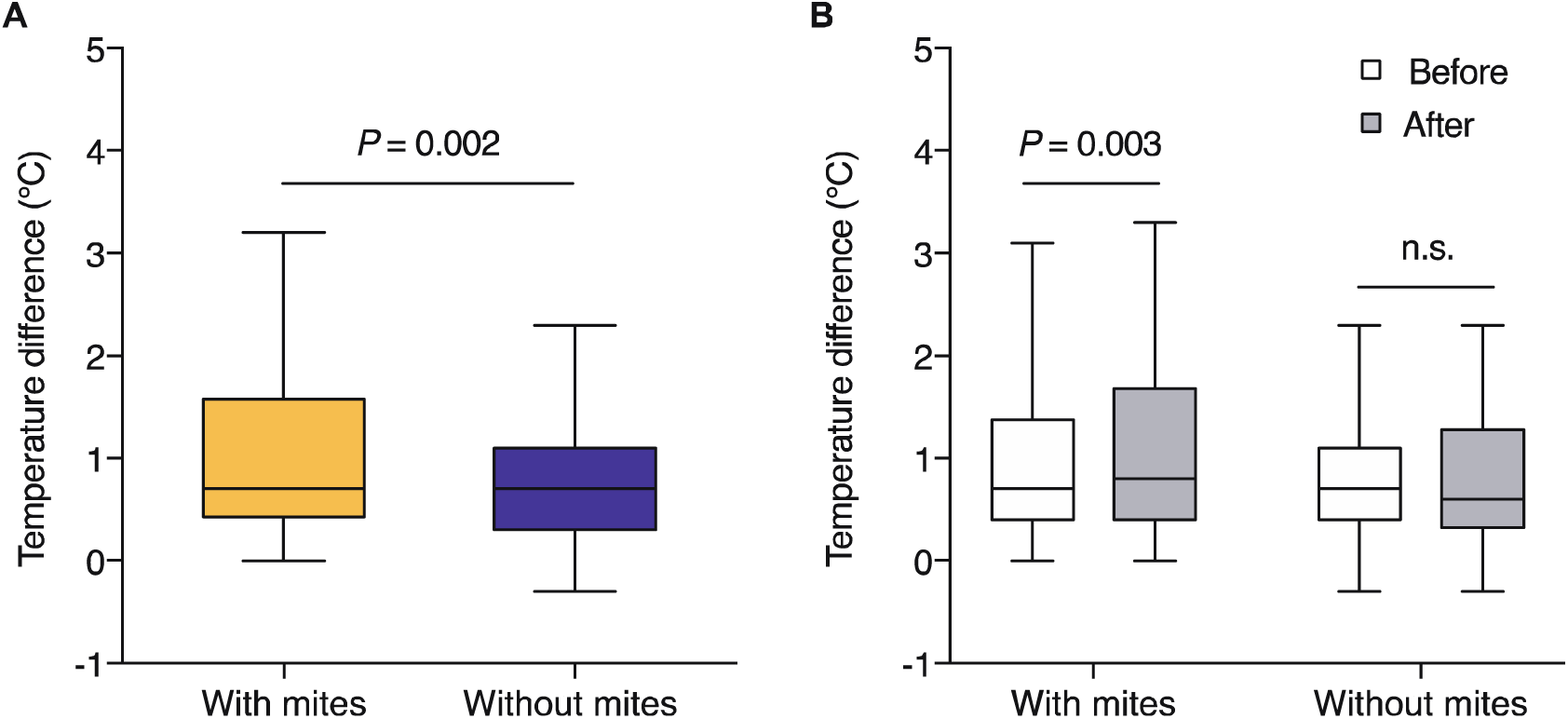
The effect of mite association on body temperature during a contest (A) and 2s before and 2s after each contest (B). Median value, inter-quartile range, and the range of data are shown in the boxplots. Each box within a plot represents data from 80 acts of aggression.

We next determined whether a raised body temperature was sufficient to improve success at winning contests, independent of an association with mites. We followed the same protocol as before, staging contests over a dead mouse between rival females that were matched in size (see Supplementary Materials). This time, neither female carried mites. Instead, prior to a contest, one of the females was placed in an incubator at 21.5°C for 30 min. This increased body temperature by 1.15± 0.14°C compared to the rival female who was not heated. The elevated body temperature increased the likelihood of this female winning the contest (*n* = 20 contests; prop.test, *P* = 0.044), with a success rate that was very similar to that induced by mites (Fig. 1). In 20 further contests, rival females each bore mites but one was cooled beforehand by placing her in an incubator at 18.5°C for 30 min. This reduced body temperature by 1.07 ± 0.76°C compared to the rival uncooled beetle. Experimental cooling also reduced the probability of winning a contest (prop.test, *P =* 0.044; Fig. 1), even though the losing female bore mites. We conclude from these experiments that mites enhance burying beetle competitive prowess by raising the beetle’s body temperature; the presence of mites alone is not sufficient to guarantee victory in a fight.

How many mites are required to raise beetle temperature sufficiently to win a contest? In natural populations, there is considerable variation in the number of mites carried by individual beetles, ranging from 0 to 285 per beetle (see Supplementary Materials and Fig. S2). We analysed how the mite density on a burying beetle influenced its body temperature by adding different numbers of mites: 0, 10, 20, or 30 mites (91% beetles carry 0-30 mites in natural populations; see Fig. S3A). We found a non-linear relationship between mite density and beetle body temperature, with a beetle’s temperature rising sharply when it carried more than 20 mites (GLMM, χ^2^ = 92.81, d.f. = 3, *P* < 0.001; Fig. S4). Adding 30 mites caused a rise in temperature that was similar to that induced by the incubator in the previous experiment (GLM, χ^2^ = 0.67, d.f. = 1, *P* = 0.414).

Next, we determined whether the thermal benefits conferred by mites differed with beetle body size (see Supplementary Materials). Even without mites, we found that larger beetles were warmer than smaller beetles (GLMM, body size effect: χ^2^ = 20.18, d.f. = 1, *P* < 0.001), which might be due to their relatively smaller surface area-volume ratio (SA/V), and consequently lower expected rates of heat loss than smaller beetles (*21*). Their consistently greater body temperature could explain, in part, why larger beetles so frequently win fights with conspecifics. The effect of the mites on beetle body temperature also varied with beetle size (GLMM, body size x mite number interaction: χ^2^ = 47.52, d.f. = 6, *P* < 0.001). Mites caused a proportionally greater increase in body temperature in smaller beetles than in larger beetles (Fig. S5), especially when 30 mites were added to the beetle (GLMM, mite number effect: χ^2^ = 30.81, d.f. = 2, *P* < 0.001).

Mites thus warm smaller beetles to a greater extent than larger beetles – but how? To determine whether mites themselves were generating heat, we compared the body temperature of freshly-killed beetles with and without mites (see Supplementary Materials). We could not detect a difference in temperature between the two treatments, (GLMM, presence v absence of mites: χ^2^ = 1.73, d.f. = 1, *P =* 0.188), suggesting that mites were not a source of heat themselves. Next, we tested whether mites cause beetles to generate heat, because they add to the mass borne by a beetle and increase the work involved in beetle locomotion. We also analysed whether mites could act as an insulator and slow down the rate at which heat generated by beetles is lost. To test these ideas, we induced small and large female beetles to walk on a motorised treadmill (see Supplementary Material and movie S2) whilst loaded with either 30 mites, or a weight equivalent to the mass of 30 mites, or bearing no load at all. Each beetle was tested with all three treatments, applied in random order across beetles. After one minute of walking on the treadmill, beetles were allowed to rest for 3 min. We measured body temperature every 10 s and 20 s during the walking and resting phases, respectively (see Supplementary Materials).

The treadmill experiments revealed that the presence of mites raised beetle body temperature during locomotion and also insulated beetles to prevent heat loss – but each of these effects was only observed when mites were carried by smaller beetles (walking: GLMM, beetle size x loading treatment x time interaction: χ^2^ = 21.36, d.f. = 12, *P =* 0.045; resting: GLMM, beetle size x loading treatment x time interaction: χ^2^ = 32.98, d.f. = 18, *P =* 0.017). Small beetles carrying mites, or weights of equivalent mass, attained a higher body temperature during locomotion than control beetles (Fig. 3A and Table S1), but there was no equivalent effect on the body temperature of large beetles (GLMM, loading treatment x time interaction: χ^2^ = 6.17, d.f. = 12, *P =* 0.907; Table S1). This is because the body temperature of larger beetles rose to a similar extent during walking on the treadmill, whether or not they were carrying anything (Fig. 3B). During the resting period, small beetles maintained a stable elevated temperature for longer when they carried mites than when they either bore a weight or were unencumbered (Fig. 3C and Table S1). By contrast, large beetles were able to maintain an elevated body temperature after locomotion without the addition of mites or weights (Fig. 3D and Table S1). These size-related effects probably arise because 30 mites add proportionally greater mass to a small beetle than to a large beetle (see Supplementary Materials). Locomotion by smaller beetles correspondingly requires more power and elevates body temperature to a greater extent (*22*). Similarly, 30 mites cover a greater proportion of a small beetle’s surface area (see Supplementary Materials and Fig. S6A), and also decrease to a greater extent the opportunity for heat loss through exposed body surface area (see Supplementary Materials and Fig. S6B). Thus, mites are more effective at reducing the rate of temperature loss on smaller individuals, but the thermal effects of mites on smaller beetles arise as a by-product of riding as passengers on the beetle, and probably did not evolve specifically to assist burying beetles.

**Fig. 3.**
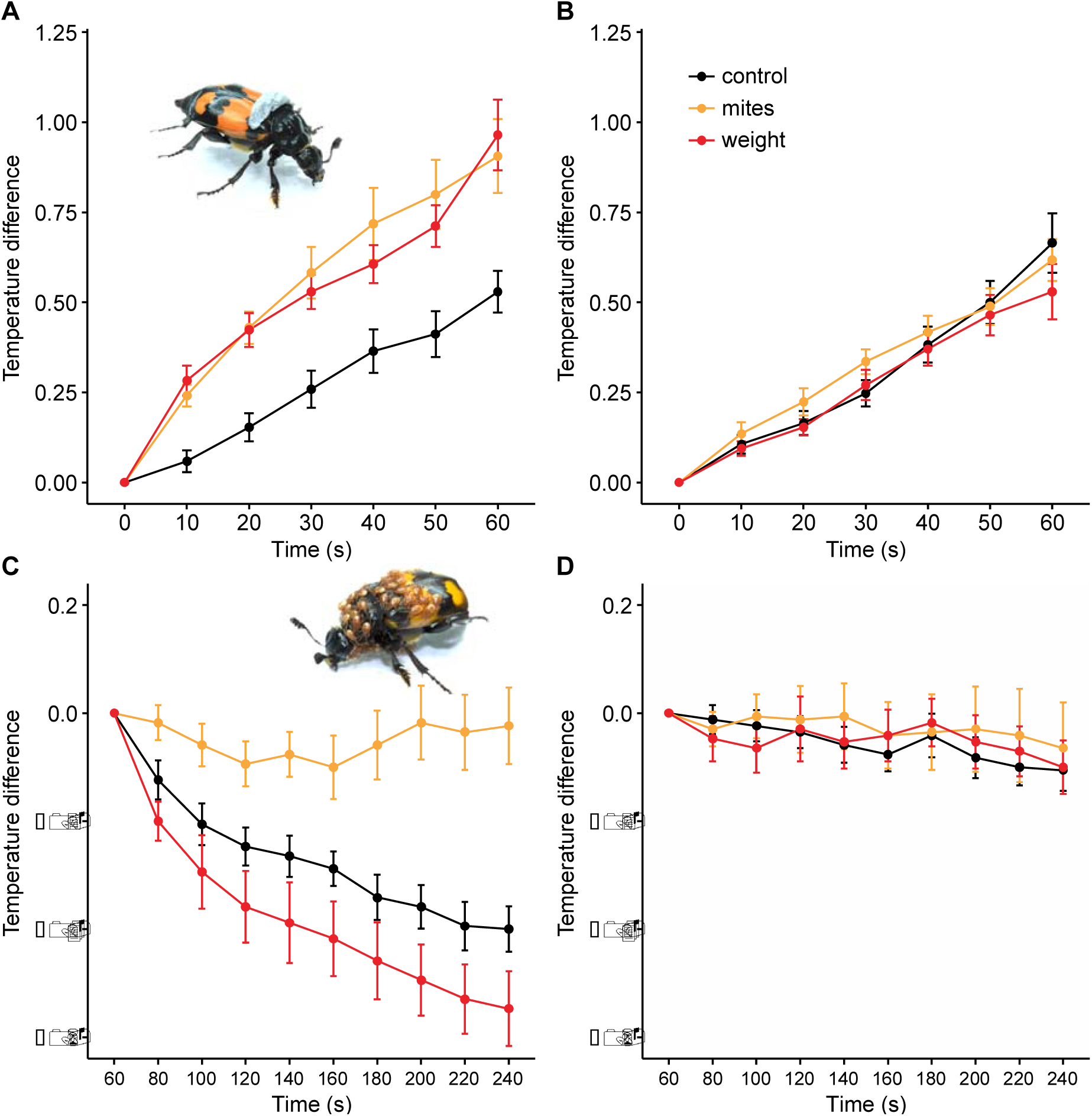
The effect of mites on burying beetle body temperature during exercise and rest. Temperature difference between loading treatments across time during walking for (**A**) small beetles and (**B**) large beetles. Temperature difference between loading treatments across time during resting for (**C**) small beetles and (**D**) large beetles. Treatments with the experimental weight (**A**) and 30 mites (**C**) are shown as insets. *n* = 17 for both small and large beetles.

Next, we determined whether the mite-induced thermal effects on smaller beetles are sufficient to compensate for the size-related disadvantage they face during contests for a carcass. We pitted focal beetles against rival medium-sized beetles (4.67 ± 0.10 mm) in contests over a dead mouse (see Supplementary Materials). Focal beetles were either small (4.29 ± 0.12 mm) or large (5.14 ± 0.16 mm) and were either loaded with 30 mites or bore no mites at all. Overall, we found that mites increased the likelihood that a smaller beetle would win the contest, but mites had no equivalent effect on larger beetles (GLM, mite x beetle size interaction χ^2^ = 4.03, d.f. = 1, *P =* 0.045; Fig. 4A). Small beetles with mites were almost three times more likely to win a contest over a carcass than were small beetles without mites (GLM, presence versus absence of mites: χ^2^ = 5.01, d.f. = 1, *P =* 0.025; Fig. 4A). Large beetles were highly successful at winning contests even without mites: bearing mites did not change their chance of victory (GLM, presence versus absence of mites: χ^2^ = 0.32, d.f. = 1, *P =* 0.574; Fig. 4A). Smaller ‘loser’ beetles thus gain more from interacting with mites than larger ‘winner’ beetles. Thus, social interactions within species determine the magnitude of the fitness benefit conferred by a second species.

**Fig. 4.**
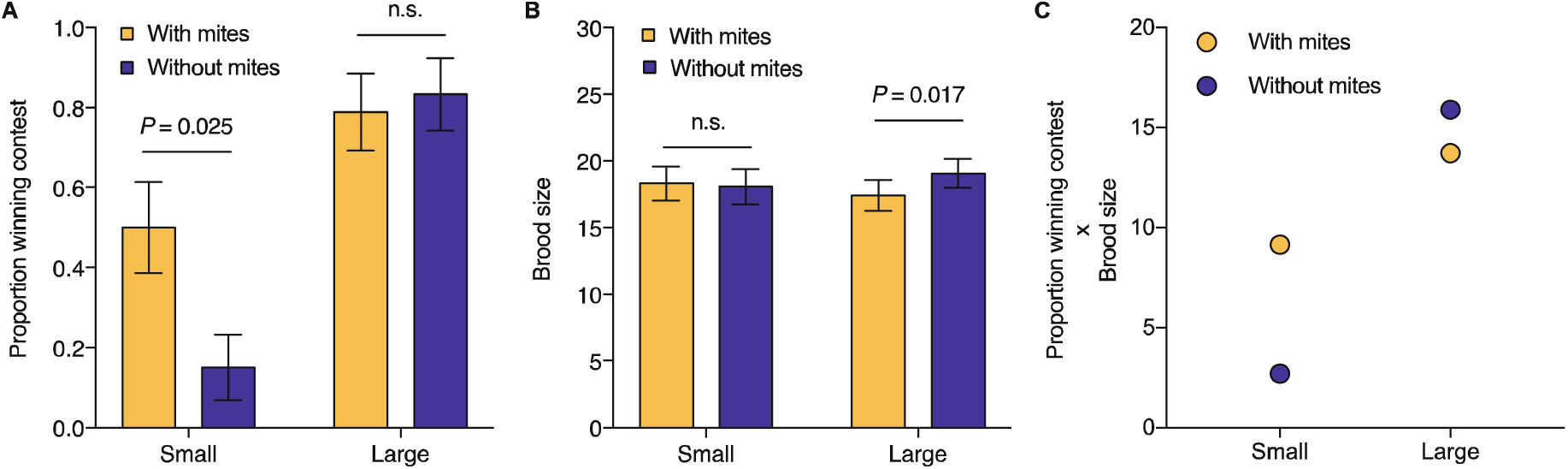
Social interactions within burying beetles determine where the effect of mites lies on a mutualism-parasitism continuum. (**A**) Proportion winning a contest when challenged with a medium-sized female for small females (*n* = 40) and large females (*n* = 37) in the presence or absence of mites. (**B**) Brood size for small females and large females when breeding in the presence or absence of mites. *n* = 34 and 33 for small females breeding with and without mites, respectively, and *n* = 33 and 28 for large females breeding with and without mites, respectively. Error bars represent mean ± SE. (**C**) Mean burying beetle fitness, calculated as the product of the mean probability of winning a contest (from **A**) and the mean number of larvae produced (from **B**), in relation to beetle size and presence/absence of 30 mites.

Having secured a carcass, beetles and mites breed alongside each other using the same carrion resource. We analysed the effect of the mites on burying beetle reproductive success by giving beetles a mouse to breed upon, uncontested (see Supplementary Materials). We compared the number of larvae produced by small and large female beetles that carried either 30 mites or carried no mites at the onset of reproduction. The effect of mites differed with beetle size (GLMM, beetle size x mite treatment χ^2^ = 4.27, d.f. = 1, *P =* 0.039; Fig. 4B). Mites reduced the brood size of large beetles (GLMM, presence versus absence of mites: χ^2^ = 5.7, d.f. = 1, *P =* 0.017; Fig. 4B) but they had no equivalent effect on small beetles (GLMM, presence versus absence of mites: χ^2^ = 0.09, d.f. = 1, *P =* 0.763; Fig. 4B). We used these experimental data to calculate the net effect of mites on the fitness of large and small beetles using the number of larvae produced as a measure of fitness. We multiplied the probability that the female would obtain a carcass in a contest (using data shown in Fig. 4A) by the mean number of larvae she would produce when breeding either with 30 mites or with no mites at all (using data in Fig. 4B). The calculations revealed that on average, 30 mites enhance the fitness of small ‘loser’ female beetles whereas they reduce the fitness of large ‘winner’ females (Fig. 4C). Social interactions within burying beetles thus define a class of individuals for whom mites are mutualists, and a distinct class for whom the mites are parasites.

In a final experiment, we considered the mites’ perspective. Whether their host beetle wins or loses a contest over carrion, mites have secured a ride to a breeding resource and can disembark to start reproduction. The limiting factor from the mites’ perspective is not the contest between beetles, but likely the relative density of conspecific mites. Previous work has shown that at high mite densities, the majority of deutonymphs do not moult into adults after acquiring a carcass, and therefore do not breed (*23*). We carried out a similar manipulation of mite density (see Supplementary Materials) and measured the probability that mites moulted in each treatment. (GLM, number of mites: χ^2^ = 104.54, d.f. = 2, *P <* 0.001; Fig. S7). We found that mites were most likely to moult when carried in groups of ten, whereas groups of thirty were least likely to moult into adults. The optimal mite density from the mites’ perspective is thus substantially lower than the optimal mite density from the perspective of a small, subordinate beetle. Competition among mites for breeding opportunities thus aligns the evolutionary interests of each individual mite away from the evolutionary interests of small beetles. This might explain why beetles are more commonly found carrying 10 mites in natural populations, and less frequently bear 30 mites (see Supplementary Material and Fig. S2).

In conclusion, our experiments show that mites and beetles are sometimes in a shared-benefit by-product mutualism, in which they work together to secure a resource they both require for reproduction (*4*). However, the extent of this interspecific mutualism is density-dependent for both species. In different ways, competitive interactions within burying beetles and within mites critically determine the fitness benefits that can be gained from interacting with the other species. Conspecifics thus play a key role in determining when mutualisms between species will persist and when they are likely to degrade into more antagonistic interactions.

## Acknowledgments

We thank the Department of Archaeology and Anthropology, University of Cambridge for the loan of a FLIR T460 infrared camera. We are grateful to C. Miller and N. Davies for their comments that improved the manuscript. We are also grateful to Sheng-Feng Shen for discussions about using thermal imaging cameras to study burying beetles. S.-J.S. was supported by the Taiwan Cambridge Scholarship from the Cambridge Commonwealth, European & International Trust. N.P.C.H. was supported by a Leverhulme Early Career Fellowship. R.M.K. was supported by a European Research Council Consolidator’s grant 301785 BALDWINIAN_BEETLES and a Wolfson Merit Award from the Royal Society. S.-J.S. and R.M.K. conceived the study. All authors designed the experiments and wrote the manuscript. S.- J.S. conducted the experiments and carried out data analysis. The authors declare no conflicts of interest. The data are deposited in Dryad.

## Supplementary Materials

### Materials and Methods

#### Study species

Burying beetles *Nicrophorus vespilloides* (hereafter simply ‘beetles’) use small carrion to breed upon, such as a dead mouse or songbird. They prepare the dead body for their own reproduction by removing the fur or feathers and rolling the carcass into a ball (*24*), which they bury below ground. The larvae hatch from eggs laid in the soil nearby, and crawl to the carcass. There they take up residence on the edible carrion nest, where they feed themselves and are fed by their parents. Parents leave the breeding attempt between 1 and 5 days after hatching, flying off to seek new reproductive opportunities (*25*). Roughly eight days after the parents first located the carcass, larvae start to disperse away into the surrounding soil to pupate. Two to three weeks later, an adult emerges and two weeks after that it attains sexual maturity.

Beetles transport mites to breed on the carrion. There are about 14 species of mites associated with burying beetle species, belonging to four different families associated with *Nicrophorus* beetles (*26*). *Poechilochirus carabi* (*Arachnia: Acari*) is the most common of these mite species (around 95% of the mites found on beetles in nature are *P. carabi* (*27*)). It exists as a species complex, consisting of races that are each specialized on a different *Nicrophorus* species (*28*). Multiple races can be found on one beetle (*28*) but they cannot be distinguished morphologically. Our unpublished behavioural data suggest that the mites used in these experiments (hereafter simply ‘mites’) comprised a mixture of the *N. humator* and *N. vespilloides* races.

*P. carabi* mites travel as deutonymphs on the burying beetles (Fig. S1). Once carrion has been located, they disembark from the beetle, moult into adults in the soil next to the carcass, and start breeding by using resources on the carrion. The presence of a carcass is essential for mites to both moult and breed successfully. The next generation of deutonymphs mostly disperses with the adult beetles as they fly off after reproduction (*29*).

All beetles and mites used in the experiments were descendants from a field-caught population collected in Madingley Woods, near Cambridge, UK (Latitude: 52.22658°; Longitude: 0.04303°) between August and October 2016. They were brought into the lab, in the Department of Zoology at the University of Cambridge, UK, and maintained on a 16:8 light:dark cycle at 20°C.

#### Burying beetle husbandry

To breed burying beetles, pairs of unrelated males and females were placed in plastic breeding boxes 2-3 weeks after eclosion (17 x 12 x 6 cm filled with 2 cm depth of moist soil), and provided with a mouse carcass (bought commercially from Livefood UK Ltd). The box was placed in a cupboard to simulate underground conditions. Eight days after pairing, as larvae were starting to disperse away from the carcass, we collected the larvae and transferred them to eclosion boxes (10 × 10 × 2 cm, 25 compartments) filled with damp soil, with one larva occupying each cell. Approximately three weeks later, after eclosion, each beetle was placed by itself in a plastic container with soil (12 × 8 × 2 cm) and fed with beef mince twice a week until it was used in the experiments.

#### Mite husbandry

To provide a continual source of mites for our experiments we started by providing pairs of burying beetles with a mouse carcass in a breeding box, as described above. Fifteen mite deutonymphs, randomly drawn from different individual beetles collected in different traps, were introduced to breed alongside the beetles (*n* = 10 breeding pairs). At the end of each breeding event, the next generation of mite deutonymphs was collected as it dispersed on beetle adults, using CO_2_ anaesthetisation. Once separated, mites were kept alongside an adult beetle in a breeding box, fed with beef mince twice a week. We replenished the mite population each month by again placing lab-bred mites with breeding pairs of beetles (*n* = 10 breeding pairs) and allowing mites to breed..

#### Natural populations

To determine the distribution of *Nicrophorus vespilloides* body size in nature, and the number of mites typically carried by each individual, we trapped beetles between June and October in 2017, which covers the entire breeding season at our study site. Traps, baited with either mice or chick carcasses, were set up constantly during this time in Gamlingay (Latitude: 52.15555°; Longitude: −0.19286°), Waresley (Latitude: 52.17487°; Longitude: −0.17354°), and Madingley (Latitude: 52.22658°; Longitude: 0.04303°) Woods in Cambridgeshire, UK. They were all checked every two weeks. Beetles were collected and brought back to the lab in the Department of Zoology. There they were anaesthetised with carbon dioxide prior body measurements and mite removal. The body size of every beetle was recorded by measuring pronotum width to the nearest 0.01 mm (this is a standard way to measure adult beetle size – see also (*30*)) and the total number of mites on each beetle was also recorded.

In total, **1369 live beetles were trapped.** The size distribution of these beetles is shown in Fig. S2. This was used to define beetles as ‘small’, ‘medium’ and ‘large’, for use in our experiments. Note that we have previously shown that the heritability of burying beetle body size does not differ significantly from 0 (*30*). Instead, variation in body size depends on the extent to which larvae are nourished during their development (*31*). Hence body size is effectively reset environmentally at each generation, and selection against smaller individuals is not sufficient to cause the evolution of increased body size within a population. From Fig. S2, *n* = 340 for small beetles (pronotum width <4.43 mm), and *n* = 360 for large beetles (pronotum width >5.08 mm).

Wild-caught beetles differed in the number of mites they carried, but only 124 beetles (9%), carried more than 30 mites (dashed line; Fig. S3A; see below for why this number is relevant). Large beetles generally carried more mites than small beetles (GLMM, χ^2^ = 29.06, d.f. = 1, *P <* 0.001; Fig. S3B). Mean number of mites carried by small and large beetles respectively were 8.09 ± 0.56 and 14.39 ± 1.32. Among these beetles, nineteen (5.59%) smaller beetles carried 30 mites or more, whereas 47 larger beetles (13.06%) carried 30 mites or more. Thus, larger beetles were more likely than smaller beetles to bear loads of at least 30 mites (GLMM, χ^2^ = 10.18, d.f. = 1, *P =* 0.001).

### Experiments

#### Effect of mites and beetle body temperature on contests between rival females

##### Treatments

We staged contests between beetles in three different ways: (i) a beetle with mites vs. a beetle without mites (*n* = 23 contests); (ii) a beetle without mites vs. a beetle without mites warmed up (*n* = 20 contests) to simulate beetle body temperature if mites were present (‘warmed’ beetle) and (iii) a beetle with mites vs. a beetle with mites cooled down (*n* = 20 contests) to simulate body temperature if mites were not present (‘cooled’ beetle). In all contests, the contestants were two females, 2-3 weeks post-eclosion, and matched in body size using measurements of their pronotum width to 0.01 mm (mean difference ± SE = 0.0095 ± 0.0010 mm). This minimized any effects of body size on contest outcome so that we could investigate effects that were due to mites and body temperature. Before the experiment females were virgins, but just prior to each contest they were mated with unrelated males, because females are typically already mated when they locate a carcass in nature. Contestants were each marked with Testors enamel paint (*32*) on the elytra before the fight, for easy identification. Each beetle was only used once, in a single contest. For beetles treated with mites, we introduced 30 mites to beetles 30 min before contests began (see below for further explanation of why we chose this mite density). Beetles that were experimentally warmed or cooled were in incubators set to 21.5°C (warmed beetles) and 18.5°C (cooled beetles), for 30 min prior to each trial. All contests took place at an ambient temperature of 20°C. Each contest was staged in a plastic container (28.5 × 13.5 × 12 cm) containing 2 cm depth of soil and a dead mouse (8-13 g). Previous studies have shown that *N. vespilloides* arriving earlier at carcasses are more likely to win any ensuing contest (*16*). Therefore, to prevent any possible confounding effects of prior ownership, the contestants were placed simultaneously in the contest arena. During each contest, individuals were able to leave the arena via a one-way valve (see (*26*) for further details).

##### Behavioural observations

A USB camera powered by a PC, with a resolution of 1920×1080 pixels, was used to record any form of aggression that occurred in the first 30 min of each trial. We classified aggressive acts as wrestling, biting, or chasing of one individual by the other (*15*). At the end of filming, we continued to observe the contest for the next 3 h, scanning the arena every 30 min to determine the outcome. When only one beetle remained on the carcass for two consecutive observations, she was deemed to be the winner.

##### Infrared thermography

Contests were also filmed with a FLIR T460 infrared camera (thermal sensitivity: <0.03°C at 30°C, 2% accuracy at 25°C) at a resolution of 320×240 pixels with frame rate at 30fps. Using the software FLIR Tools+ 6.4 (Copyright 2018 FLIR Systems, Inc; http://www.flir.com), the body temperature of beetles was measured at the centre of the thorax, whereas the temperature of soil was determined as the average temperature measured inside a 2 cm diameter circle randomly oriented adjacent to where a beetle was sitting on the soil. We determined the emissivity of beetle cuticle using methods described in the Supplemental Information of Smolka *et al.* 2012 (*33*). To ensure accurate measurement of temperature, all measurements were taken at a constant distance of 25 cm from the surface being measured. The calibrated infrared emissivity of beetles, and soil, was set to 0.95 (0.947 ± 0.02, *n* = 23); in this way all measures were scaled in relation to the thermal radiation emitted by a black body. We synchronised footage from the infrared camera with our standard film of the contest to determine beetle body temperature and soil temperature during acts of aggression, 2s before a fight started, and 2s after a fight ended.

#### Effect of mite density on beetle body temperature

We measured female body size, by measuring the width of her pronotum. We then added groups of 10 mite deutonymphs sequentially to the same individual female. Thus each female started with 0 mites, then had 10, then 20, and finally 30 mites added (*n* = 45 beetles). To control for the potential order effects of mite association, we also manipulated mite number in reverse order by initially adding 30 mites and then removing 10 mites at a time (*n* = 45 beetles). We measured beetle body temperature 30 min after the addition, or removal, of 10 mites, using the infrared camera described above (Fig. S4). The effect of mite density on beetle body temperature, in relation to beetle body size, is shown in Fig. S5.

#### Are mites a source of heat?

To determine whether changes in beetle body temperature were due to mites or beetles, we added or removed mites in batches of 10 in exactly the same way as described above (adding mites: *n* = 10 beetles; removing mites: *n* = 10 beetles), but this time using dead female beetles. The dead beetles were killed just before the experiment by exposing them to −20°C for an hour. They were then put in an incubator at 20°C until they acclimated to environmental temperature in the lab. At this point they were used in the experiment.

#### Treadmill experiments

To determine whether the beetle’s elevated body temperature was due to carrying the weight of mites and/or whether mites serve as an insulating ‘blanket,’ we continuously measured changes in beetle body temperature using infrared thermography across time for small (4.05 ± 0.18 mm, *n* = 17) and large (5.21 ± 0.12 mm, *n* = 16) beetles, as they walked on a motorised treadmill modified from a lab tube rotator (Movie S2). Beetles walked on the continuously moving track for one minute. The track was set to move at a constant speed of 4.7cms^-1^ (see Movie S2), which is the slowest walking speed of a beetle carrying no mites (mean speed, 6.74 ± 1.72 cms^-1^; *n* = 13 beetles). This period of exercise was followed by a three minute rest period. Beetle body temperature was measured at the centre of the thorax every 10s and 20s during walking and resting phases, respectively. We tested each beetle three times, randomly mixing up the sequence in which they carried the following loads: (i) control – no load, (ii) 30 mites (7.78 ± 1.79 mg), and (iii) an experimental weight, equivalent to 30 mites (8.04 ± 0.59 mg). The experimental weight consisted of a blob of Blu-Tack ^®^ that was gently moulded and attached to the front portion of the elytra, which is where *P. carabi* mites are typically located (Fig. S1). This allowed us to temporarily manipulate the body mass of beetles without causing trauma. Mean body masses for small and large beetles respectively were 120.99 ± 16.17 mg and 222.44 ± 15.96 mg. Thus, carrying mites increased the body mass of small and large beetles by 6.5% and 3.6% respectively (*t* test, *t* = 5.33, *P* < 0.001), whereas carrying experimental weights increased the body mass of small and large beetles by 6.7% and 3.7% respectively (*t* test, *t* = 8.11, *P* < 0.001).

#### Measurement of beetle surface area, surface area covered by mites, and surface area-volume <u>ratio</u>

To further understand how mites were able to insulate beetles, we estimated a beetle’s surface area, the area covered by different numbers of mites, and the surface area:volume ratio, of small (4.08 ± 0.33 mm, *n* = 11) and large (5.21 ± 0.14 mm, *n* = 10) beetles. We added groups of 10 mites, up to a total of 30 mites, to the same individual female, or else added 30 mites, and then removed 10 mites at a time, to control for the order effects of manipulation. After addition or removal of mites, we allowed 10s for the mites to freely distribute themselves over the surface of the beetle. The ventral and dorsal surface of every beetle was then photographed next to a scalebar, at a constant distance and under the same lighting conditions. We similarly took photos of beetles carrying an experimental weight equivalent to 30 mites (see details in Treadmill experiments section). All digital images were analysed using image analysis software (ImageJ, https://imagej.nih.gov/ij/). We extracted data from calibrated images by calculating the area of the dorsal and ventral surfaces, the area covered by 0, 10, 20 or 30 mites, and the area covered by the experimental weight. For simplicity, we assumed that beetles are two-dimensional, meaning we could estimate total body surface area as the sum of the area of the dorsal and ventral views. After photography, beetles were euthanised by exposing them to −20°C for an hour, all mites were removed, and the volume of each beetle was determined by the water displacement method. Since we were interested in understanding how the presence of mites influenced the potential for heat loss, we calculated surface area:volume ratios as the ratio of uninsulated beetle surface area (sum of the beetle surface area – sum of the surface area of *n* mites) divided by beetle volume.

We began by checking that our methods were not confounded by only taking measurements from the dorsal and ventral surfaces of beetles. This is because some mites were present on the lateral surfaces of beetles, but they were not included in our estimation of the beetle surface area covered by mites. To determine whether or not this was a potential confounding effect, we counted the number of mites that attached to the sides of beetles and tested whether the number differed with the mite density on the beetle (10, 30, 30 mites), the beetle’s size (small/large), and the interaction between mite treatment and beetle size. Neither the interaction term (χ^2^ = 2.00, d.f. = 2, *P* = 0.368), nor beetle size (χ^2^ = 1.54, d.f. = 1, *P* = 0.214) significantly influenced the number of mites that were found on the lateral surfaces of beetles, although we did find more beetles on lateral surfaces at greater mite densities (χ^2^ = 15.74, d.f. = 2, *P* < 0.001). This means we have under-estimated the surface area covered by mites, particularly at high mite densities. Therefore, the results of the analyses that follow are biased to be conservative.

We found that 30 mites covered a greater proportion of the surface area of a small beetle than they did on a large beetle (22.3% and 8.8% respectively; GLMM, beetle size x mite number interaction: χ^2^ = 75.59, d.f. = 3, *P <* 0.001; Fig. S6A). Furthermore, relative to the beetle’s volume, 30 mites covered a greater surface area on small beetles than did any other mite load, leaving a much smaller area exposed and therefore uninsulated (GLMM, beetle size x mite number interaction: χ^2^ = 35.93, d.f. = 4, *P <* 0.001; Fig. S6B). These results explain why we found a non-linear relationship between mite number and beetle body temperature (Fig. S5).

By contrast, the experimental weight had no insulative properties (Fig. 3C), because it covered a much smaller surface area. Nevertheless, it occupied proportionately more surface area on a small beetle than on a large beetle (small beetles: t-ratio = 14.25, *P* < 0.001; large beetles: t-ratio = 3.34, *P* = 0.030; Fig. S6A). For large beetles, whether beetles carried 30 mites, or the experimental weight, or no mites at all, there was no significant change in the surface area that was exposed, relative to the beetle’s volume (30 mites: t-ratio = 2.54, *P* = 0.266; experimental weight: t-ratio = 1.71, *P* = 0.787; Fig. S6B). This may explain why rate of heat loss in all three treatment groups for large beetles was essentially the same (Fig. 3D).

#### Effect of mites on contests between beetles that differed in size

We again staged contests between two females over a dead mouse, but this time we ensured that the contestants differed in size. In one treatment, focal beetles were ‘small’ (4.29 ± 0.12 mm, *n* = 40), while in a second treatment focal beetles were ‘large’ (5.14 ± 0.16 mm, *n* = 37). In each contest, focal beetles were pitted against a different medium-sized beetle (4.67 ± 0.10 mm; Fig. S2). In roughly half the contests, the focal beetle bore 30 mites, which were added beforehand as described above (*n* = 20 small beetles, *n* = 19 large beetles). In all other details, the procedure for staging the contest was exactly as described above, except that we did not film these contests.

#### Effect of mites on burying beetle reproductive success, with respect to beetle body size

We determined the effects of mites on reproductive success by breeding small (4.20 ± 0.19 mm, *n* = 67) and large (5.17 ± 0.17 mm, *n* = 61) females with either 0 or 30 mites on 8-13 g mouse carcasses (9.51 ± 1.05 g, *n* = 128). For the treatment with mites, 30 deutonymphs were added as we introduced females to breeding carcasses. These beetles did not experience a contest prior to breeding. At dispersal, we counted and weighed all dispersing third-instar larvae as a proxy of breeding success.

#### Effect of mite density on mite reproductive success

To investigate how mite density influences mite deutonymphs’ moulting rate, we repeated the experiments described in (*23*). Mites were kept as groups of 1 (*n* = 19), 2 (*n* = 22), 4 (*n* = 22), 8 (*n* = 18), 10 (*n* = 20), 12 (*n* = 20), 16 (*n* = 19), 20 (*n* = 19), or 30 (*n* = 20), on pieces of moist filter paper within Petri dishes (diameter 60 mm, height 15 mm). Each group was provided with a piece of lamb liver (0.6-0.8 g) to trigger moulting. After 2 days, we checked the number of deutonymphs that moulted into males or females or remained unmoulted.

### Statistical analyses

All analyses were conducted in R version 3.4.3 (*34*). Generalized linear mixed models (GLMM) were used in the package *lme4* (*35*) with fixed and random factors to analyse the effects of mite number on body temperature and the contest outcomes.

#### Effect of mites and beetle body temperature on contests between rival females

To examine the contest outcomes of each of the three experimental treatments (i.e. with mites vs. without mites, without mites/warmed vs. without mites, and with mites/cooled vs. with mites), we performed the proportional test using the *prop.test* function. This analysis determines whether the outcome is significantly biased away from the chance outcome of 50% of contests being won by the focal beetle. To investigate mite effects on body temperature during each aggression, we included mite treatment, carcass mass, and relative difference in body size, i.e. [(focal female pronotum width – nonfocal female pronotum width)/focal female pronotum width] as fixed factors. Each contest consisted of a single outcome (win or lose), but within each contest multiple aggressive acts could be recorded for each beetle. Therefore, we included the ID of each trial as a random factor. Since we had no *a priori* expectation as to how mites or temperature should affect beetle aggressive behaviour, we grouped all behaviour types (wrestling, biting, or chasing) together in our analyses, while still recording the total number of aggressive acts occurring within a contest. For each act of aggression that occurred between beetles within a contest, we also analysed temperature changes 2s before and 2s after the act of aggression for beetles with and without mites by including timing (before/after), carcass mass, and relative body size difference as fixed factors, and order of the aggressive act (i.e. first aggressive act, second aggressive act, and so on) nested within trial ID as a random factor. We included ‘block’ as a fixed factor, since the experiment was carried out with beetles from two consecutive generations.

#### Effect of mite density on beetle body temperature

To examine how mite number affected body temperature, we included the difference between body temperature and soil temperature as a dependent variable; mite number treatment (0, 10, 20, and 30 mites as a categorical variable), sequence of mite association (increase or decrease), and body size of each individual as fixed factors, and individual ID as a random factor.

To further understand the effects of mites on the proportional increase in temperature across body size, temperature difference and the ratio of temperature difference (dividing temperature difference with mites by temperature difference without mites) were included as dependent variables. Mite number, body size, and their interaction, and order of mite association were included as fixed factors, and individual ID as a random factor.

#### Are mites a source of heat?

To examine whether mites themselves generate heat, we analysed the difference between body temperature and soil temperature for ten freshly-killed beetles in the same way as described above.

#### Treadmill experiments

To assess whether the equivalent weight of mites differentially affects the body temperature for small and large beetles as they walk on a treadmill, we analysed the interacting effects between treatments (control, mite, and weight) and body size (small or large) across the walking period (0-60s) on the temperature difference (focal body temperature – body temperature at time 0s). Focal body temperature was sampled every 10s. To assess whether mites provide an insulative ‘blanket effect’ to reduce beetles’ heat loss while resting after walking, we analysed the interacting effects between treatments (control, mite, and weight), body size (small or large) and time (60-240s) on the temperature difference (focal body temperature – body temperature at time 60s). Focal body temperature was sampled every 20s. For both analyses, individual ID was included as a random factor since individual beetles were repeatedly sampled.

#### Measurement of beetle surface area, surface area covered by mites, and surface area:volume ratio

We analysed whether carrying mites or the experimental weight changed the proportion of surface area exposed, and whether this differed between small beetles and large beetles. In this analysis, the dependent variable was the proportion of beetle surface area covered by mites (or the experimental weight), calculated as the sum of the surface area covered /sum of the dorsal and ventral surface areas. This measure was log-transformed prior to analysis to meet assumptions of data normality. We included the load borne by the beetle (i.e. 10, 20 or 30 mites, or the experimental weight) and body size (small or large) as fixed factors, and also the interaction between them. Individual ID was included as a random factor since individuals were repeatedly sampled. We used a similar model to determine how these variables influenced a beetle’s surface area, relative to its volume. The only difference was that in this model the dependent variable was calculated as the (sum of the beetle surface area – sum the surface area covered by mites)/beetle volume.

#### Effect of mites on contests between beetles that differed in size

We analysed the effects of mites on the outcome of a contest when beetles differed in body size using a binomial distribution. The outcome (winner or loser) was included as a dependent variable, while mite treatment, body size (small or large), and carcass mass were included as fixed factors.

#### Effect of mites on burying beetle reproductive success, with respect to beetle body size

We analysed the effects of mites on brood size at dispersal, for small and large beetles, by using a Poisson distribution and including mite treatment and carcass mass as independent variables.

#### Effect of mite density on mite reproductive success

To test for an influence of mite density on the moulting rate of mites, we analysed the effect of number of mites as a continuous predictor on the proportion of moulted mites in a GLM using a binomial distribution with a logit link function. The model that fitted best was a second-order polynomial regression (χ^2^ = 101.59, d.f. = 1, *P <* 0.001; Fig. S7), replicating the findings of a previous study (*23*). We also checked whether there was an effect of the amount of liver that was provided to the mites on the likelihood of moulting. Liver mass had no effect on proportion of mites moulting (χ^2^ = 0.93, d.f. = 1, *P =* 0.335).

**Figure S1.**
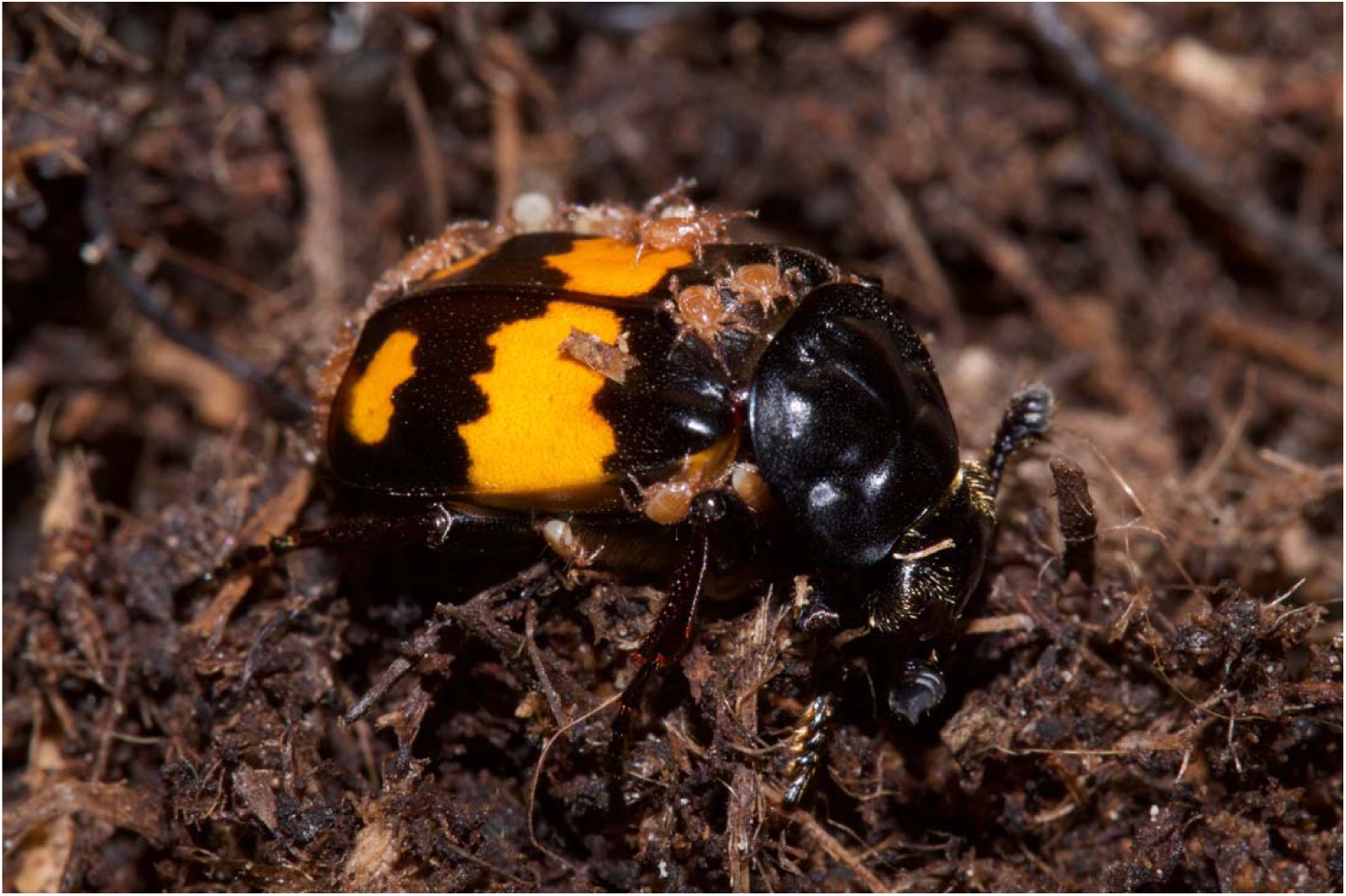
A burying beetle *N. vespilloides* bearing mites from the *P. carabi* species complex. Photo by S.-J. Sun.

**Figure S2.**
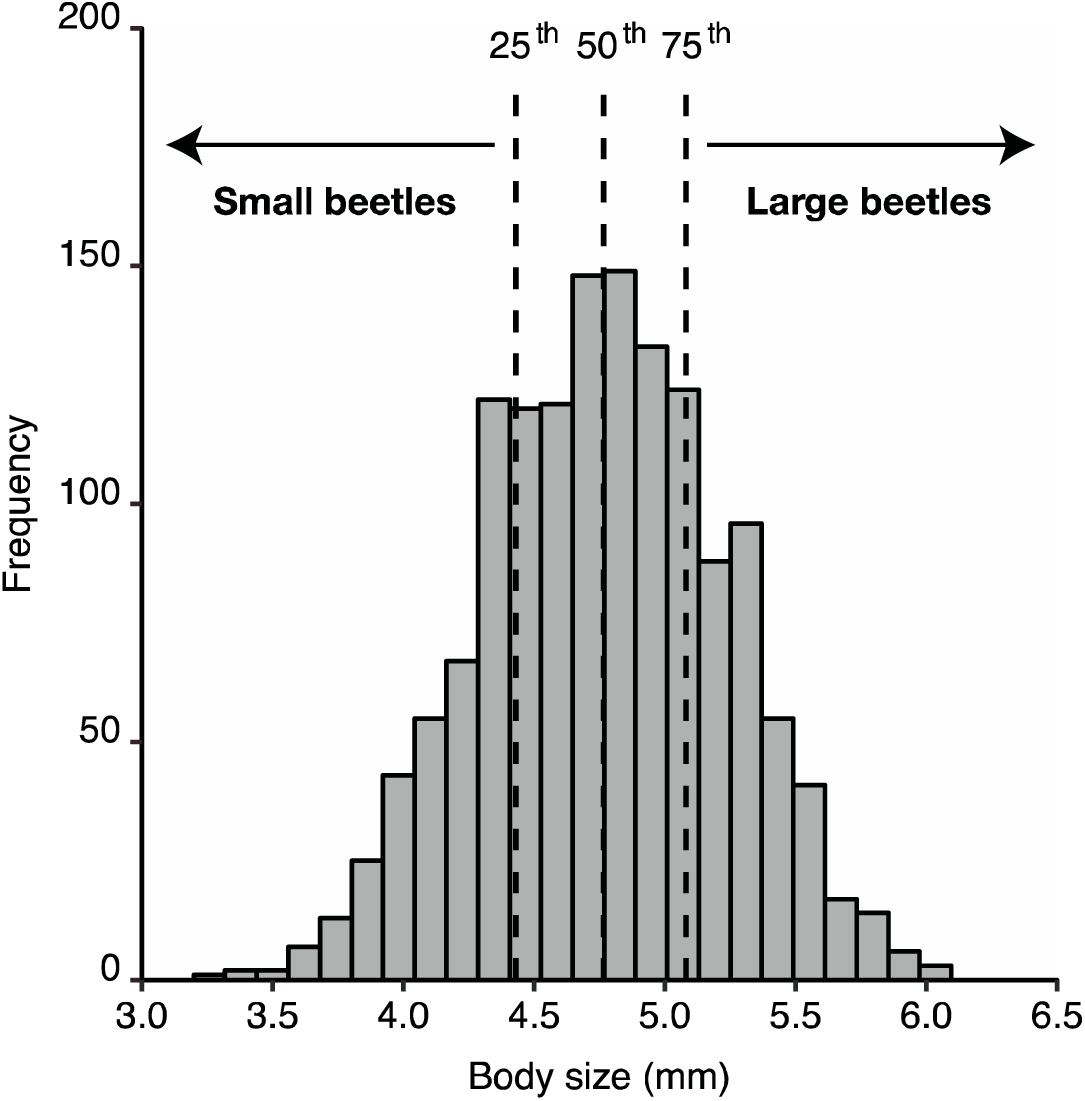
Frequency distribution of the body size, given by pronotum width, of wild-caught *N. vespilloides* beetles across three woodlands in Cambridgeshire. *n* = 1452 beetles. Dashed lines indicate 25^th^, 50^th^ and 75^th^ percentiles. Beetles falling below the 25^th^ percentile i.e. < 4.43mm pronotum width were defined as ‘small’ in our experiments, whereas beetles above the 75^th^ percentile i.e. >5.08mm pronotum width were defined as ‘large’.

**Figure S3.**
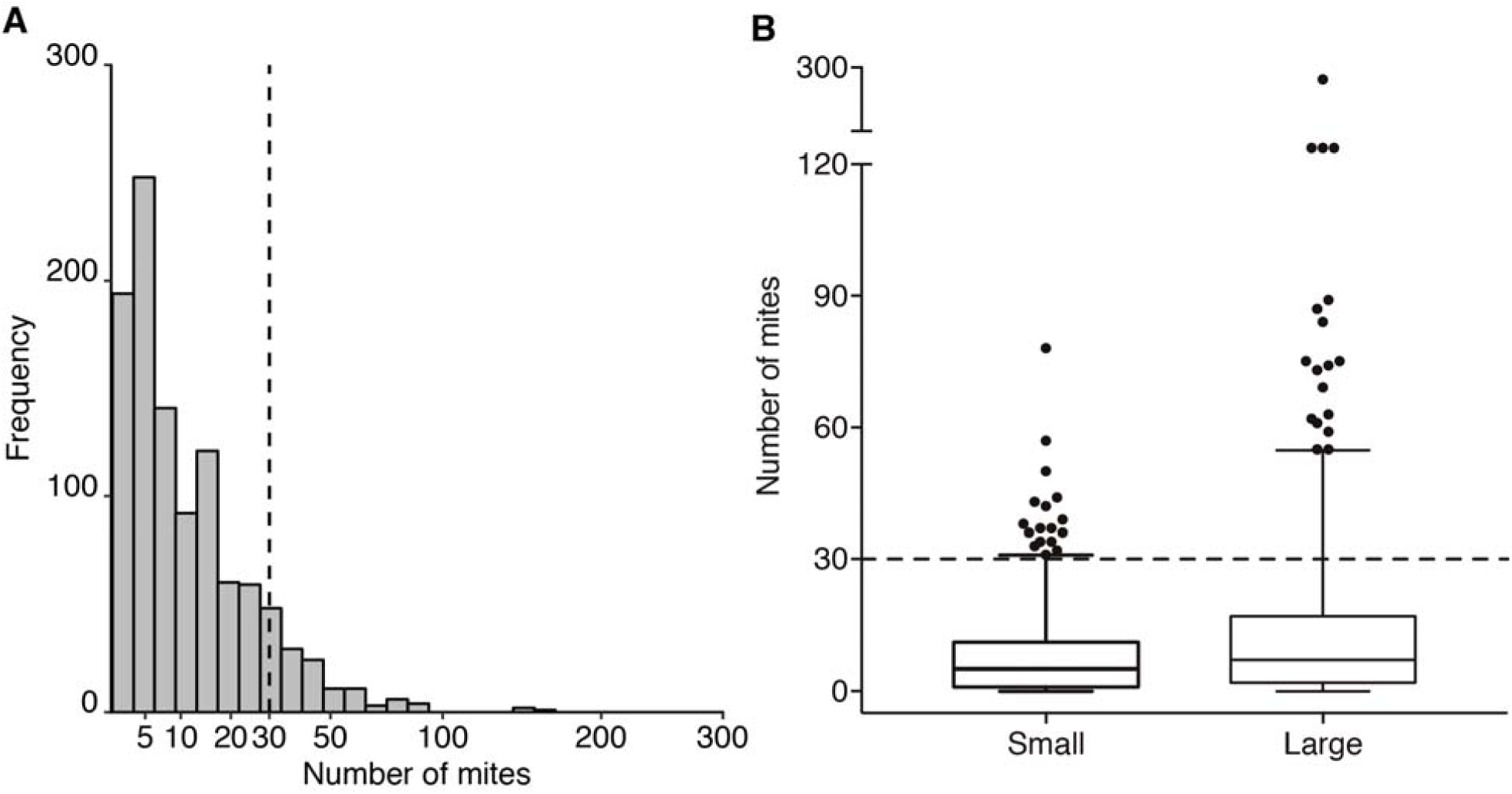
Frequency distribution of the number of mites carried, for all wild-caught *N. vespilloides* (A), and the number of mites carried by small and large wild-caught *N. vespilloides* (B). (A) Frequency distribution of the number of mites carried by beetles across three woodlands in Cambridgeshire, UK – the source of the beetles used in our experiments. *n* = 1369 live beetles. (B) Large beetles (*n* = 360) carried more mites compared to small beetles (*n* = 340). Median value, inter-quartile range, and 5^th^ and 95^th^ percentile (error bars) are shown in the boxplots. Outliers are shown as points. The dashed lines in both figures indicate the number of individuals naturally carrying 30 mites, which is the maximum mite load used in the experiments.

**Figure S4.**
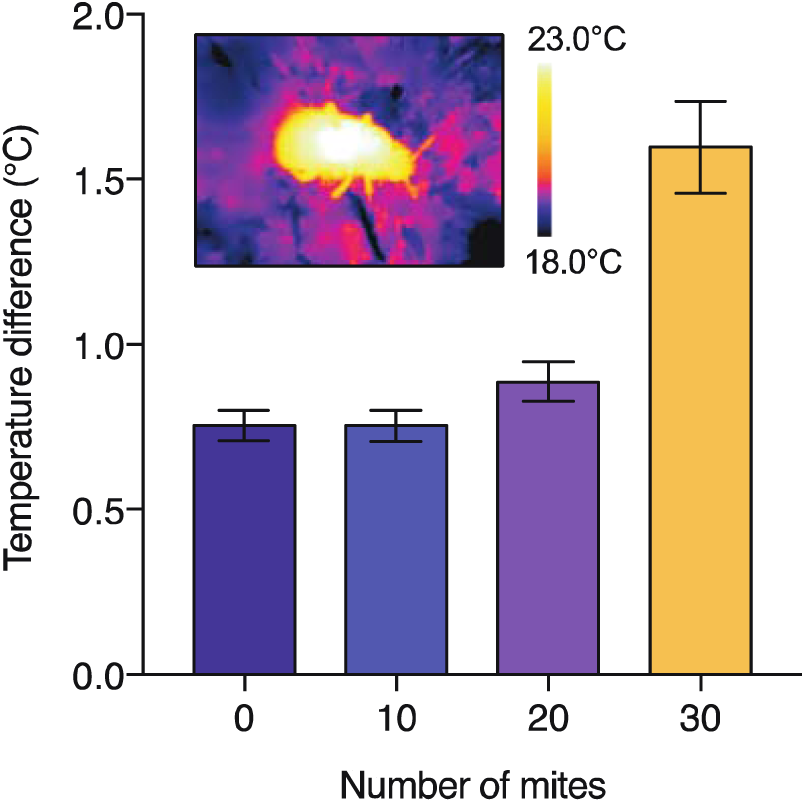
The relationship between mite load and beetle body temperature, relative to soil temperature. *n* = 90 beetles, each subjected to all four mite treatments. Inset is a thermal image of a burying beetle carrying mites. Means ± SE are shown.

**Figure S5.**
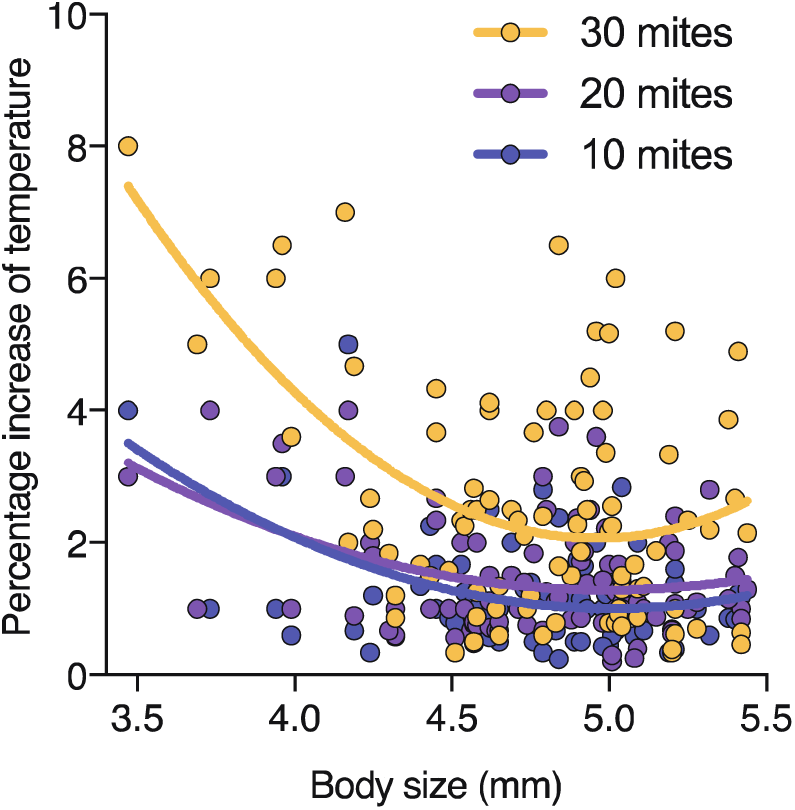
The proportional increase in body temperature as a consequence of carrying mites in relation to body size, given by pronotum width. Solid lines are regression lines fitted in GLMM. Temperature difference refers to the difference between beetle body temperature and the temperature of the soil on which the beetle is sitting.

**Figure S6.**
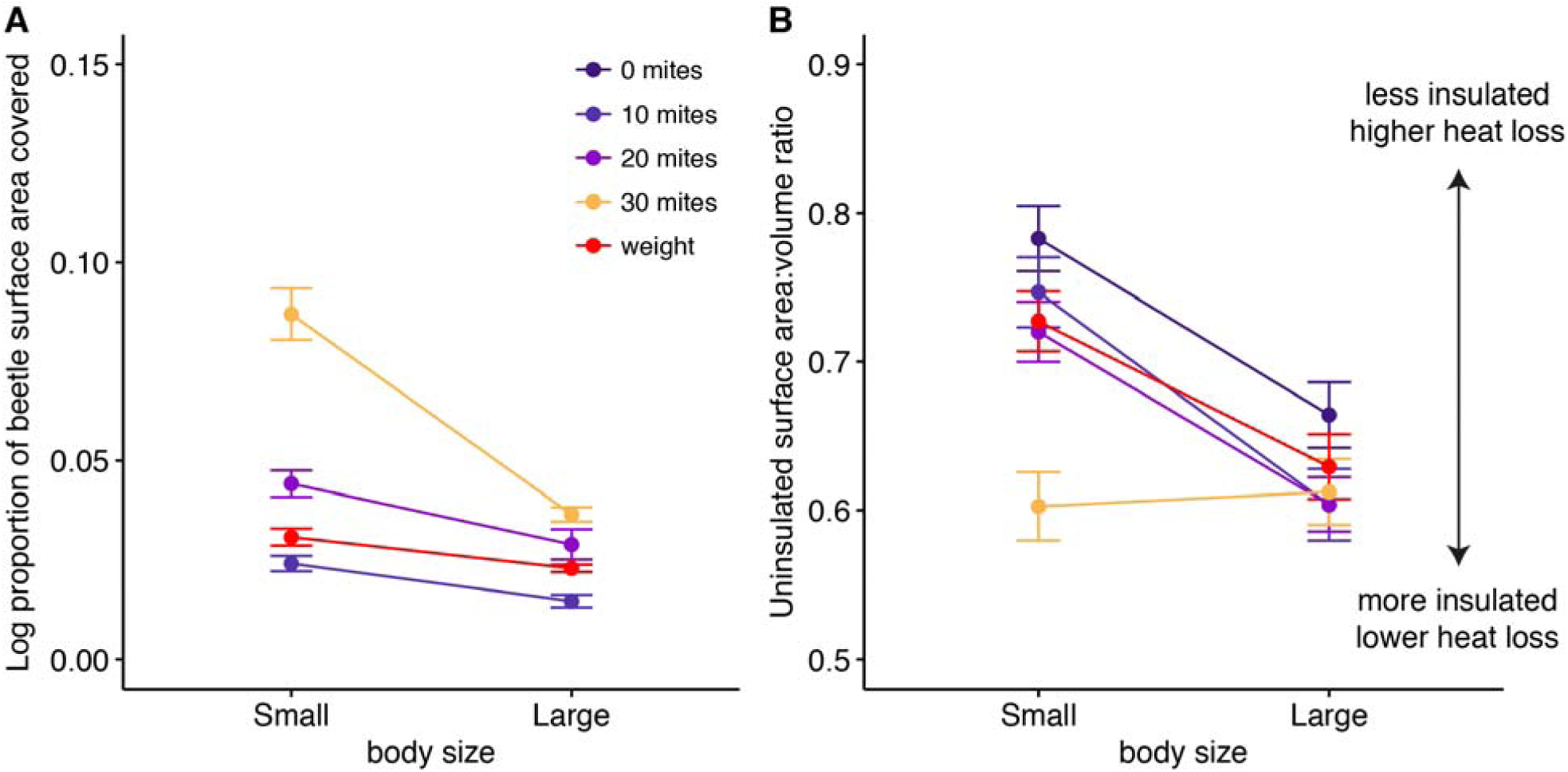
Effect of mite density on the proportion of beetle surface area covered (A) and the size of the uninsulated area that remains, relative to the beetle’s volume (B). Values for the experimental control weight, used in the treadmill experiments, is shown for comparison in both figures. (**A**) 30 mites cover more surface area on a small beetle than on a large beetle. Note also that the experimental weight has the same mass as 30 mites but covers a much smaller surface area (**B**) The size of the uninsulated area on a burying beetle, relative to its volume, is markedly lower for smaller beetles carrying 30 mites compared to similarly sized beetles carrying fewer mites. *n* = 11 and 10 for small and large beetles, respectively.

**Figure S7.**
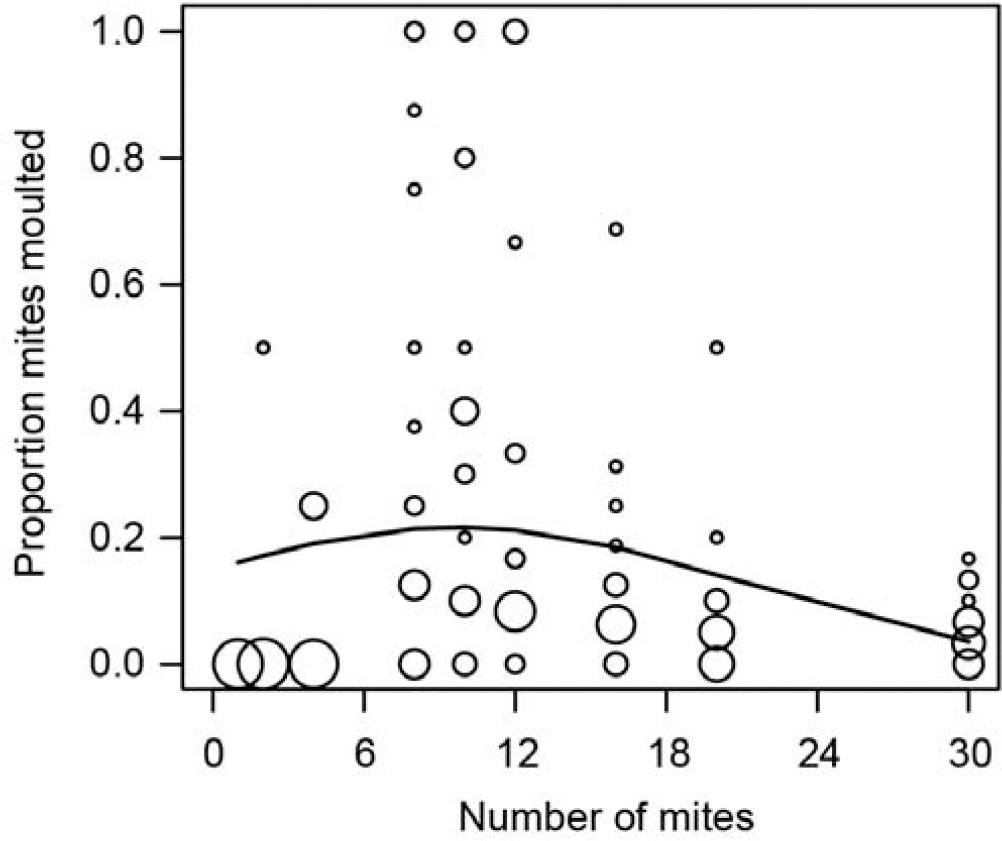
Effect of mite density on mite reproductive success. The proportion of mites that moulted upon reaching a carcass, in relation to mite group size (*n* = 179). Circles denote different groups of mites on a single beetle, and circle size increases with the number of congruent data points. The curved line denotes a statistically significant relationship, as predicted from a GLM.

**Table S1.** Results from the models analysing changes in body temperature as a function of the load carried by beetles during the treadmill experiments. The dependent variables were the temperature difference (walking: focal body temperature – body temperature at time 0s; resting: focal body temperature – body temperature at time 60s). Loading treatments (control, mite, and weight) were included as independent variables and beetle identity was included as a random factor. Significant factors are shown in bold.

| Response variables | Temperature difference (walking) |  |  | Temperature difference (resting) |  |  |
| --- | --- | --- | --- | --- | --- | --- |
| Independent variables | $\chi^2$ | d.f. | *P*-value | $\chi^2$ | d.f. | *P*-value |
| Intercept | 0.00 | 1 | 1.000 | 0.00 | 1 | 1.000 |
| Loading treatments | 0.00 | 2 | 1.000 | 0.00 | 2 | 1.000 |
| **Time** | **65.58** | **6** | **<0.001** | **68.62** | **9** | **<0.001** |
| **Loading treatments\*Time** | **23.35** | **12** | **0.025** | **61.34** | **18** | **<0.001** |

*Large beetles*
| Response variables | Temperature difference (walking) |  |  | Temperature difference (resting) |  |  |
| --- | --- | --- | --- | --- | --- | --- |
| Independent variables | $\chi^2$ | d.f. | *P*-value | $\chi^2$ | d.f. | *P*-value |
| Intercept | 0.00 | 1 | 0.963 | 0.086 | 1 | 0.770 |
| Loading treatments | 4.53 | 2 | 0.104 | 2.66 | 2 | 0.265 |
| **Time** | **462.70** | **6** | **<0.001** | 11.51 | 9 | 0.242 |

**Movie S1.** A beetle with mites attacking a beetle without mites. Inset is the corresponding thermal imaging video.

**Movie S2.** A beetle walking on a motorised treadmill.

